# Reproducibility of Diffusion, Shape, and Connectivity Metrics Across Scanners: Implications for Multi-Site Tractography

**DOI:** 10.64898/2026.04.15.718542

**Authors:** Sachit Anand, Fang-Cheng Yeh, Siva Venkadesh

## Abstract

Multi-site diffusion MRI studies face scanner-induced variability that can obscure biological signal. Harmonization methods such as ComBat have been developed to address this, but have been evaluated primarily on diffusion scalar metrics. Whether scanner reproducibility differs across fundamentally distinct tract-derived representations has not been systematically compared.

Here, we compared the reproducibility of three metric families (diffusion, shape, and connectivity) across 36 association tracts using the MASiVar dataset (5 subjects, 4 scanners, 27 sessions). We assessed intraclass correlation coefficients (ICC) and multivariate subject discrimination at baseline, under dimensionality reduction, and after ComBat harmonization.

At baseline, shape metrics showed the highest reproducibility (median ICC 0.69), followed by connectivity (0.49) and diffusion (0.34). Shape and connectivity achieved comparable subject discrimination (both 1.75), significantly exceeding diffusion (1.23). ComBat harmonization improved all families but harmonized diffusion (0.58) remained below unharmonized shape (0.69), indicating that metric family selection remains consequential even after harmonization.

Under low-dimensional representation, connectivity showed the largest gains (ICC 0.86, subject discrimination 3.0), exceeding other families at any dimensionality. Analysis of principal component loadings identified a small number of cortical regions per tract (median 6) that capture 95% of the reproducible connectivity signal, providing a per-tract reference for selecting the most informative regions in future multi-site studies.

These findings indicate that the choice of which tract-derived metrics to analyze in multi-site studies deserves at least as much consideration as how to harmonize them.

## 1. Introduction

Multi-site diffusion MRI studies increasingly rely on tract-derived measures to characterize white matter organization, yet the reproducibility and scanner sensitivity of commonly used tract representations remain incompletely understood. Prior work has demonstrated that diffusion scalars, tractography-based connectivity measures, and geometric descriptors can each exhibit substantial scanner effects, and that harmonization methods such as ComBat(Fortin et al., 2017) and RISH(Mirzaalian et al., 2016) may mitigate these effects.

However, existing evaluations of scanner variability, including large-scale harmonization efforts that have successfully established normative reference curves for diffusion scalars across tens of thousands of subjects(Cetin-Karayumak et al., 2024; Zhu et al., 2025, 2019), have focused primarily on metrics within a single family, typically FA and MD. Recent work has begun addressing connectivity directly, with(Kurokawa et al., 2021) demonstrating that site effects in structural connectomes persist after signal-level harmonization and require correction at the matrix level, and(Onicas et al., 2022) applying ComBat to connectome-derived graph metrics. However, a systematic comparison of scanner reproducibility across fundamentally different tract representations (diffusion microstructure, tract geometry, and regional connectivity) has not been undertaken, nor has prior work examined whether multivariate subject discrimination, which captures the joint signal across features, reveals reproducibility structure invisible to univariate measures such as the intraclass correlation coefficient (ICC)(Shrout and Fleiss, 1979). This matters because different metric families occupy different positions in the processing hierarchy: diffusion metrics summarize voxel-level microstructure, shape metrics capture tract-level geometry, and connectivity metrics describe region-level overlap patterns. These successive levels of spatial abstraction may confer different degrees of inherent robustness to scanner variability.

We address this gap using a subject-anchored reliability framework that decomposes total variability into subject-specific and scanner-driven components. By comparing these representations under both native and dimension-matched (PCA) conditions, and before and after ComBat harmonization, our results offer practical guidance for selecting tract-derived metrics in multi-site studies and clarify how representation dimensionality shapes reproducibility.

## 2. Methods

### 2.1 Dataset

MASiVar Cohort II(Cai et al., 2021) comprises 27 diffusion MRI sessions acquired from 5 healthy adults across 4 scanners at 3 sites. Preprocessed data were downloaded from OpenNeuro (ds003416), where all scans had been processed with PreQual v1.0.0, including Marchenko-Pastur denoising, intensity normalization, susceptibility and eddy current distortion correction, motion correction, and slice-wise signal dropout imputation.

### 2.2 Pipeline

Diffusion data were reconstructed using DSI Studio(Yeh, 2021) (version “Hou” Aug 2024) with generalized q-sampling imaging (GQI, diffusion sampling length ratio = 1.25) and resampled to 2.5 mm isotropic resolution. Volumes were rigidly aligned to AC-PC orientation prior to reconstruction. Thirty-six bilateral association tracts were identified using automatic tract recognition, which matches subject streamlines to a population-based tractography atlas via nonlinear registration and Hausdorff distance filtering.

For each tract, three families of metrics were extracted (Figure 1). Shape metrics (19 per tract) were derived from the geometric properties of reconstructed streamlines(Yeh, 2020), including tract count, mean length, span, curl, elongation, total volume, quarter-segment volumes, surface area, irregularity, and endpoint area, radius, and branch volume for each terminus. Diffusion metrics (13 per tract) were computed as mean values along each tract’s streamlines, comprising DTI-derived scalars (FA, MD, AD, RD), GQI-derived measures (QA, GFA, ISO, RDI, HA), and restricted diffusion indices (RD1, RD2, nRDI at b = 2000 and b = 4000).

**Figure 1.**
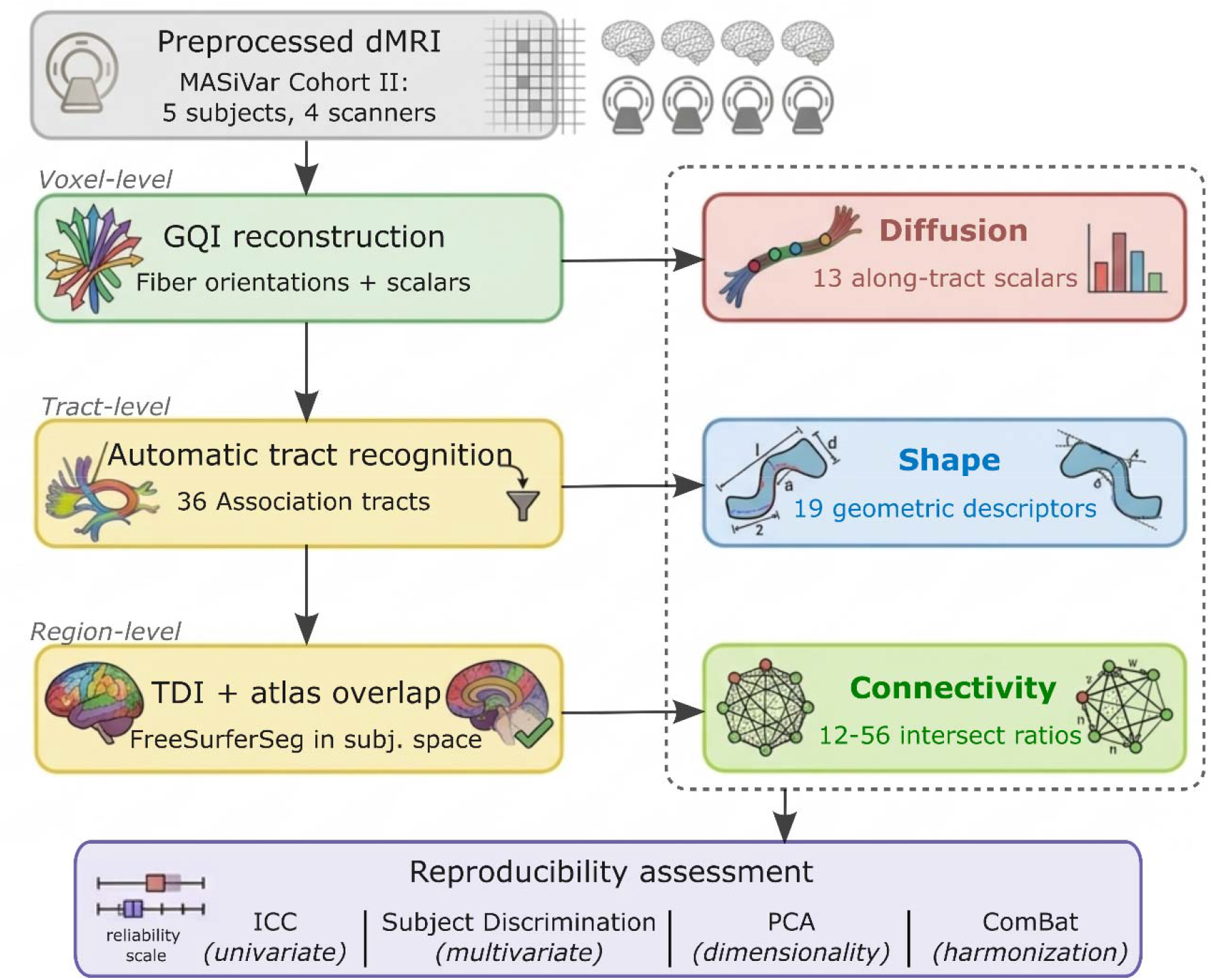
Processing pipeline and metric family overview. Preprocessed diffusion MRI data from MASiVar Cohort II (5 subjects, 4 scanners) are processed through successive stages yielding three metri families at different levels of spatial abstraction: diffusion (voxel-level), shape (tract-level), and connectivity (region-level). All three families are evaluated using four reproducibility assessments (bottom).

Connectivity metrics were derived from tract-to-region overlap analysis. Quantifying the spatial overlap between axonal pathways and parcellated regions has a long history in both tractography(Donahue et al., 2016; Hagmann et al., 2008; Owen et al., 2015) and viral tracing connectomics(Oh et al., 2014). (Yeh, 2022)defined a population-level tract-to-region connectome that quantifies connectivity for each tract independently; here, we extend it to individual subjects. For each tract, a track density image (TDI) is computed by rasterizing all streamlines onto the native diffusion voxel grid, counting distinct streamlines per voxel. The FreeSurferSeg parcellation(Fischl, 2012) is warped nonlinearly to subject space, and for each region, the intersect ratio is computed as the fraction of region voxels where tract density exceeds a noise floor (0.5% of maximum TDI value). This yields a value between 0 and 1 per tract–region pair, representing the spatial overlap between each tract and each cortical or subcortical region. The number of regions with nonzero overlap varies per tract (range: 12–56), reflecting each tract’s cortical territory.

### 2.3 Reproducibility metrics

We assess reproducibility at two complementary levels.

**ICC (univariate):** For each individual metric within a tract, the intraclass correlation coefficient (Shrout and Fleiss, 1979) quantifies the proportion of total variance attributable to true subject differences:

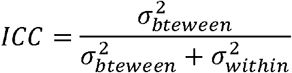

Here, 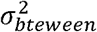 reflects biological differences across subjects and 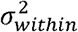 reflects measurement variability across scanners for the same subject. This evaluates whether a single measurement is stable across scanners.

#### Subject discrimination (multivariate)

Rather than evaluating metrics independently, subject discrimination assesses whether the full profile of metrics within a family jointly distinguishes individuals across scanners. For shape and diffusion, features are z-score normalized prior to distance computation to account for heterogeneous measurement scales; connectivity features, which are uniformly expressed as intersect ratios, are used without standardization to preserve the natural variance structure.

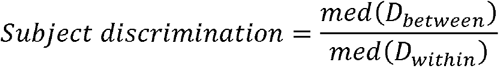

Here, *D*_*between*_ is the set of pairwise Euclidean distances between sessions from different subjects, and *D*_*within*_ is the set of pairwise Euclidean distances between sessions from the same subject on different scanners. Both are computed over all metrics in the family jointly. A value of 1 indicates no subject separability; higher values indicate that between-subject distances exceed within-subject (scanner-driven) distances.

### 2.4 Dimensionality analysis (PCA)

The three metric families differ substantially in native dimensionality: shape contributes 19 features per tract, diffusion 13, and connectivity 12–56 depending on the tract’s cortical extent. To enable fair cross-family comparison and to assess how reproducibility depends on representational dimensionality, principal component analysis (PCA) was applied independently within each metric family for each tract. For shape and diffusion, features were standardized prior to PCA to account for heterogeneous scales (correlation-based PCA); for connectivity, PCA was applied to the raw covariance matrix to preserve the natural variance hierarchy among intersect ratios. ICC and subject discrimination were recomputed at each dimensionality from 1 to 20 retained components. This analysis reveals whether reproducible subject-specific signal is concentrated in a few dominant modes or distributed across the full feature space, a distinction that proves critical for connectivity metrics in particular.

### 2.5 Harmonization (ComBat)

To evaluate the effect of scanner harmonization on reproducibility, ComBat(Fortin et al., 2017) was applied to the baseline (non-PCA) features within each metric family. Scanner identity served as the batch variable, and subject identity was included as a biological covariate to preserve inter-individual differences during harmonization. ICC was recomputed after harmonization to quantify the gain attributable to removing scanner-specific variance.

## 3. Results

### 3.1 Baseline metric family comparison

Figure 2 illustrates the raw variability across scanners and subjects for a representative tract (left arcuate fasciculus), showing how within-subject (scanner-driven) and between-subject differences co-exist in the raw metric values.

**Figure 2.**
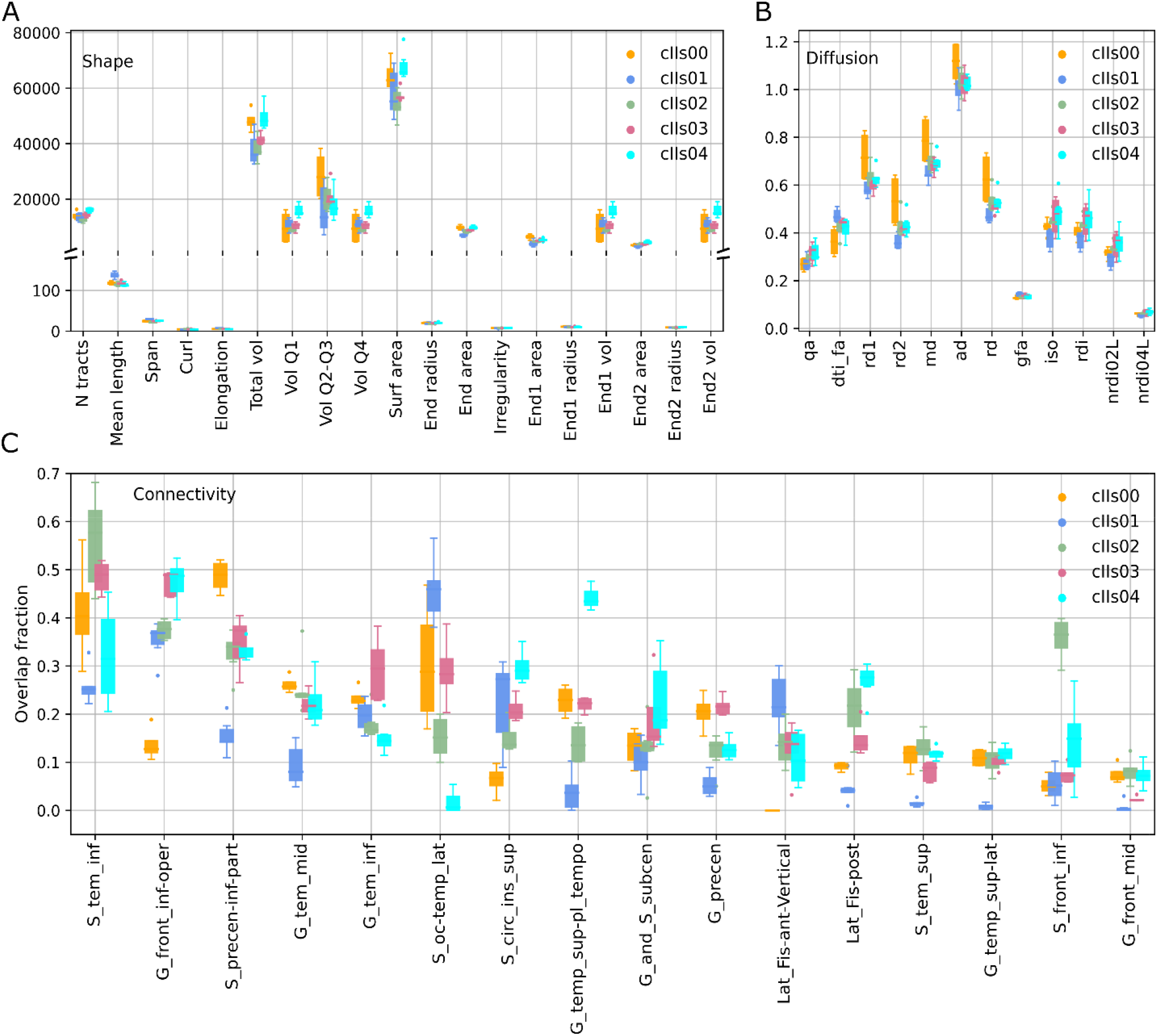
Raw variability of tract-derived metrics across scanners and subjects for the left arcuate fasciculus. Box plots show the distribution of metric values across scanning sessions for each of the 5 subjects (color-coded). (A) Shape metrics. (B) Diffusion metrics. (C) Connectivity metrics (intersect ratios for the top overlapping regions).

At baseline, shape metrics showed the highest reproducibility across the 36 association tracts (Figure 3), with a median ICC of 0.69, followed by connectivity (0.49) and diffusion (0.34). All pairwise differences were significant (Shape > Diffusion, p < 0.001; Shape > Connectivity, p < 0.001; Connectivity > Diffusion, p < 0.01). For subject discrimination, shape and connectivity were statistically indistinguishable (medians 1.75 and 1.75, respectively; p = 1.000), while both significantly exceeded diffusion (1.23; p < 0.001). Shape thus leads in per-metric reproducibility, but shape and connectivity are equally effective at distinguishing individuals when the full feature profile is considered. Diffusion is lowest on both measures.

**Figure 3.**
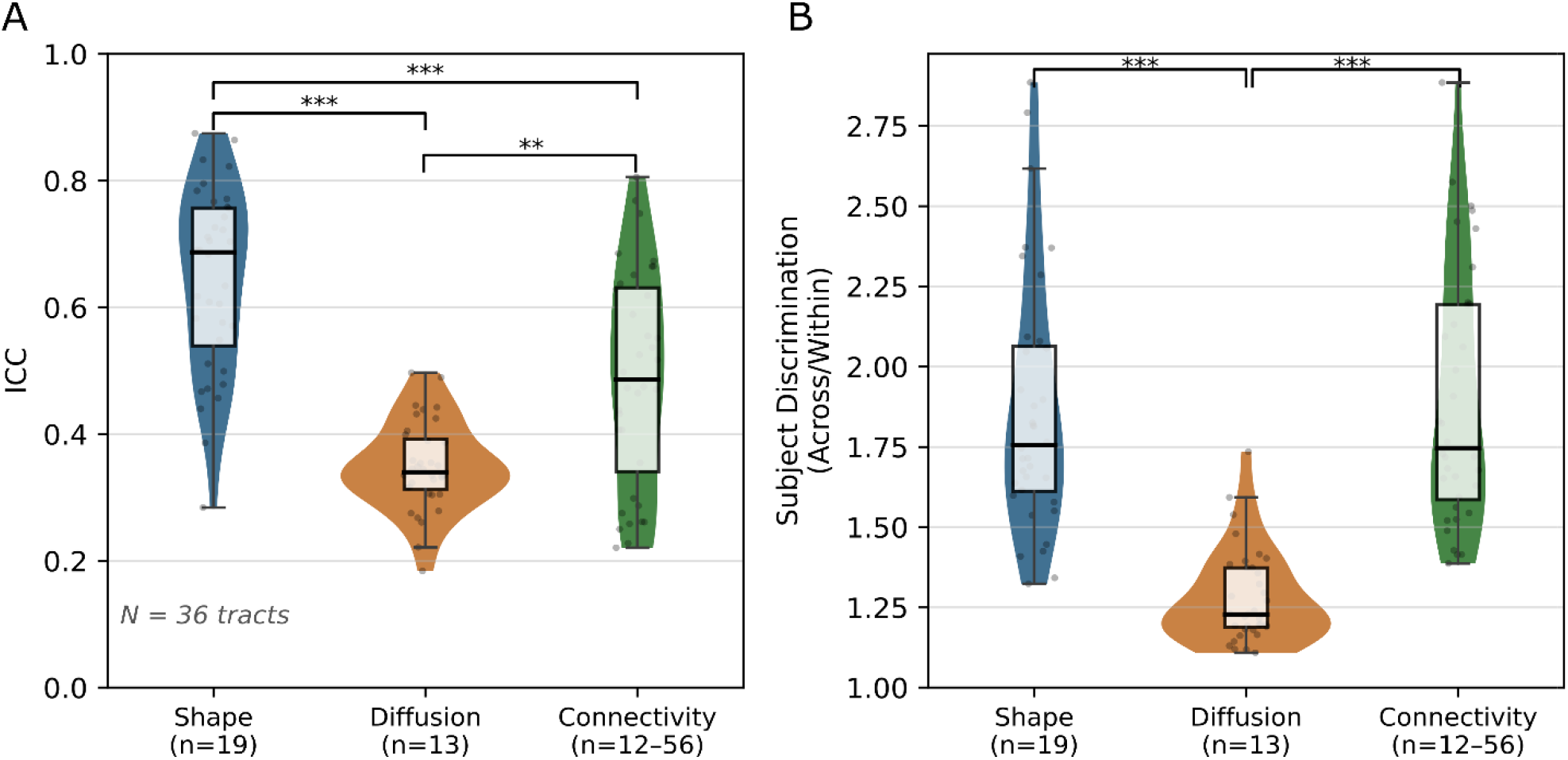
Baseline reproducibility and subject discrimination across metric families. (A) Violin plot of tract-level ICC for shape, diffusion, and connectivity across 36 association tracts. Significance brackets indicate pairwise Wilcoxon signed-rank tests. (B) Multivariate subject discrimination (ratio of across-subject to within-subject Euclidean distance) for each metric family.

ICC varied substantially across tracts within each family (Supplementary Figure S1). Longer association tracts with distinctive geometry, such as PAT-R (0.88), SLF2-R (0.86), ILF-L (0.83), and ILF-R (0.82), showed the highest shape reproducibility, likely because their size and trajectory make them reliably identifiable across scanners. The lowest-performing tracts were predominantly cingulum subdivisions, including CgL-PHP (0.28), CgL-PH (0.39), CgL-FPH (0.46), and CgL-SLF1 (0.47), which are shorter and more sensitive to registration variability. Connectivity ICC showed the most tract-to-tract variability, with some tracts matching or exceeding shape (e.g., PAT-L connectivity 0.81) while others fell well below diffusion (e.g., HA-L connectivity 0.23, EC-L connectivity 0.22), reflecting dependence on the number and distinctiveness of cortical overlap regions per tract.

### 3.2 Effect of dimensionality reduction

At one component, all pairwise family differences were significant (paired Wilcoxon, p < 10^-^□ each; Figure 4A–B. Connectivity ICC reached approximately 0.86, far above its baseline of 0.49. Shape ICC rose to approximately 0.75 (from 0.69), while diffusion showed a more modest increase to approximately 0.48 (from 0.34). Subject discrimination followed a similar pattern: connectivity reached approximately 3.0 at one component, well above its baseline of 1.75. Shape rose to approximately 2.4 and diffusion to approximately 1.4 at two components. At 1–3 components, connectivity ICC and subject discrimination exceeded all other families at any dimensionality. For subject discrimination, all families converged to their respective baseline values at high component counts, confirming that full-rank PCA recovers the native feature space. ICC does not show this convergence because it is a per-component property rather than a joint distance measure.

**Figure 4.**
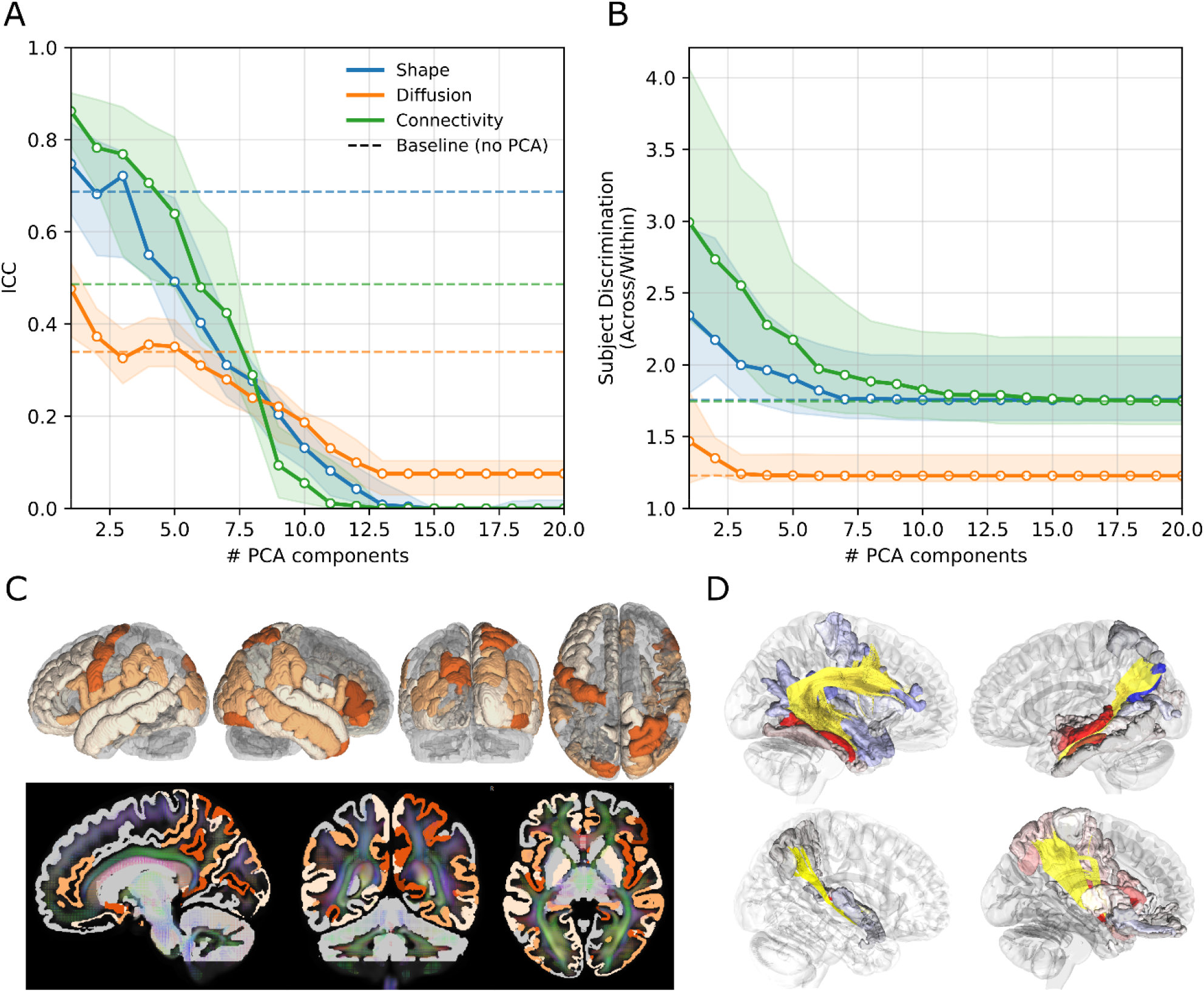
Reproducibility of low-dimensional representations across metric families. (A) Median ICC as a function of retained PCA components for shape (blue), diffusion (orange), and connectivity (green). Dashed lines indicate baseline ICC; shaded regions show interquartile range across 36 tracts. (B) Median subject discrimination as a function of PCA components. (C) Cortical regions contributing to reproducible individual variation in connectivity. Color intensity indicates the number of tracts for which a region is a principal driver (light peach = 1, deep/burnt orange = 4). Bottom row: slices including subcortical structures, overlaid on a QA map with fiber orientation coloring. (D) Example tracts illustrating signed PC1 loading patterns. Warm colors indicate positive loadings, cool colors negative; tract streamlines in yellow.

All three families showed improved reproducibility at low component counts, but connectivity benefited disproportionately due to its larger and more correlated feature space (see Discussion). Because PCA is fit per tract, the regions driving the first component differ across tracts. Inspection of PC1 loadings confirms that the highest-loading regions correspond to known termination territories for each tract (e.g., amygdala for hippocampal-amygdala tracts, insular cortex for extreme capsule, intraparietal sulcus for SLF; Supplementary Table S1). Retaining regions that account for 95% of PC1 loading variance selects a median of 6 regions per tract (range 2–14), adapting to each tract’s loading concentration. Mapping these regions onto cortical surface across all 36 tracts shows that 126 of 192 FreeSurferSeg regions contribute to reproducible individual variation in connectivity (Figure 4C).

### 3.3 Effect of harmonization

All within-family ComBat improvements were significant (paired Wilcoxon, p < 10^-^□for each family; all 36 tracts improved in every family; Figure 5). Shape increased from 0.69 to 0.77 (+12%), diffusion from 0.34 to 0.58 (+72%), and connectivity from 0.49 to 0.62 (+27%). Despite these gains, the cross-family hierarchy persisted: harmonized diffusion (0.58) remained significantly below unharmonized shape (0.69; r_rb = −0.41). Metric family selection is thus consequential even after harmonization.

**Figure 5.**
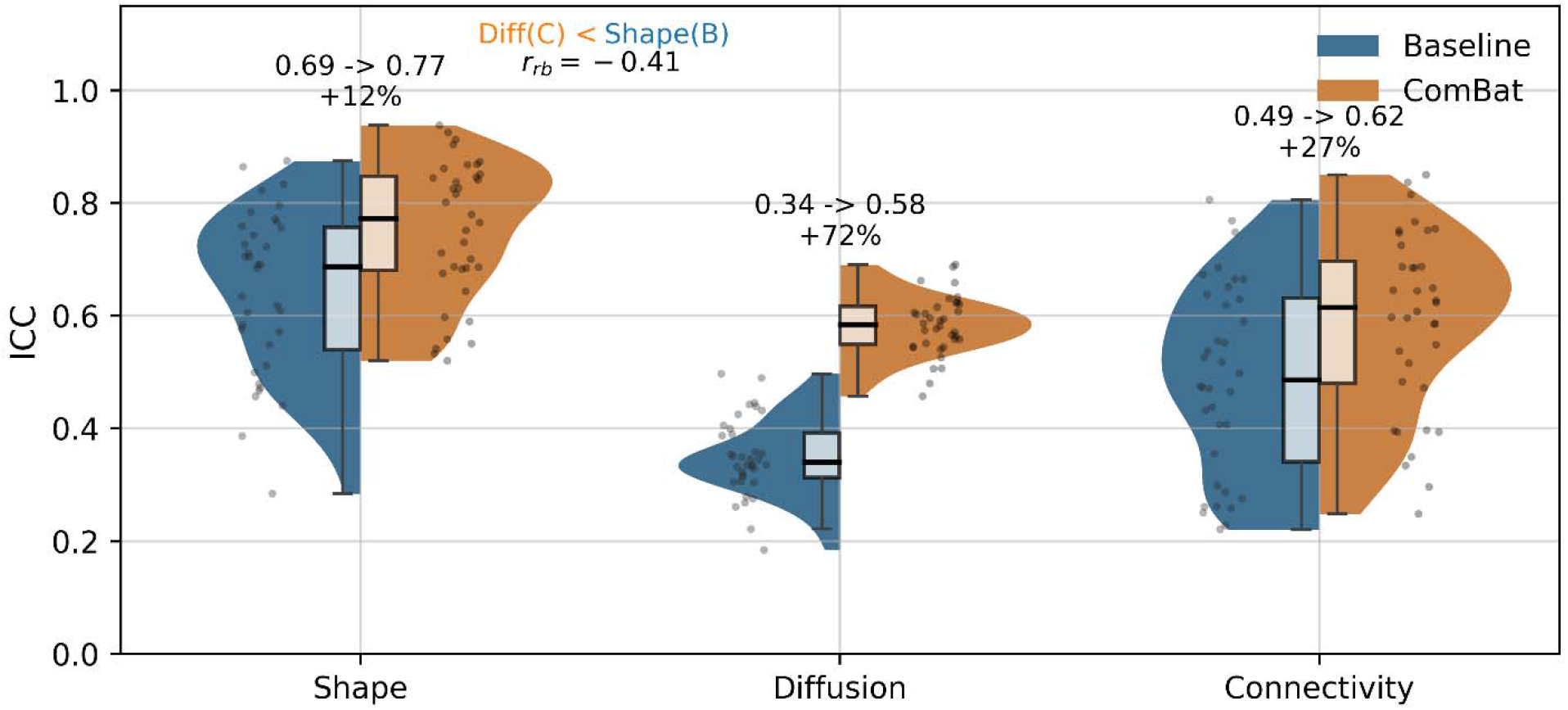
Effect of ComBat harmonization on reproducibility across metric families. Split violin plots showing the distribution of tract-level ICC before (baseline, blue) and after (ComBat, orange) harmonization for shape, diffusion, and connectivity. Individual points represent tract-specific ICC values. Annotations indicate median ICC, percent change, and the rank-biserial correlation (r_rb) comparing harmonized diffusion to unharmonized shape.

It is worth noting that for connectivity, dimensionality reduction (ICC 0.86 at one component) produced larger gains than ComBat harmonization (ICC 0.62), whereas for shape and diffusion the two approaches yielded comparable improvements. This pattern suggests that for connectivity, appropriate representation of the distributed biological signal yields larger reproducibility gains than scanner correction alone.

## 4. Discussion

Diffusion MRI tractography has become a central tool for investigating white matter organization across clinical and developmental applications(Assaf et al., 2019). As multi-site consortia scale these studies, the choice of which tract-derived measures to analyze becomes consequential, yet is rarely treated as a methodological variable. In practice, many studies prioritize diffusion tensor–derived scalars such as FA and MD, which are among the most established metrics in the literature(Figley et al., 2021; Jones et al., 2013; Nir et al., 2017). This prevalence is understandable: DTI scalars are straightforward to compute and have accumulated extensive normative data, and the field’s harmonization infrastructure has developed primarily around them(Cetin-Karayumak et al., 2024; Zhu et al., 2025, 2019). Our results suggest, however, that this concentration may not reflect an optimal allocation of effort: diffusion metrics are the least reproducible across scanners among the three families we evaluated, and this disadvantage persists even after harmonization.

### 4.1 Why shape metrics lead at baseline

Shape metrics describe macroscopic tract geometry derived from reconstructed streamlines. Each step in the processing chain, from voxel-level signal to fiber orientation to streamline trajectory to geometric summary, integrates over progressively larger spatial scales and reduces sensitivity to voxel-level noise and some scanner-dependent variability. Properties such as volume, length, and curvature reflect aggregate characteristics of many (correlated) streamlines rather than signal intensity at any individual voxel, making them less sensitive to differences in gradient strength, noise floor, or signal-to-noise ratio. While connectivity overlap is also derived from streamline trajectories, it remains more sensitive to endpoint definitions and spatial alignment, which may limit the extent of this robustness in individual connections. This geometric stability also underlies shape’s advantage in ICC: individual differences in tract anatomy, such as the size and curvature of major association pathways, are well captured by shape descriptors, exhibit between-subject variability that exceeds measurement noise, and remain relatively consistent across scanning conditions. Notably, connectivity achieves comparable subject discrimination despite lower per-metric ICC, suggesting that its biological signal is distributed across many features rather than concentrated in individual metrics.

### 4.2 Why connectivity uniquely benefits from PCA

All three metric families contain internally correlated features: diffusion scalars covary through the shared tensor, shape metrics scale together with tract size, and connectivity overlap values reflect shared spatial patterns. PCA exploits this redundancy in each family, explaining why even shape and diffusion ICC improve at low component counts. Connectivity benefits disproportionately, however, because its subject-specific signal is distributed across many small, consistent regional differences that aggregate into a strong first component. The result is a representation where 1–3 components per tract capture nearly all reproducible variance, an effective subject fingerprint inaccessible from any single connectivity value.

Examination of PC1 loadings supports the anatomical validity of this compression. For each tract, the highest-loading regions correspond to expected termination zones, and the top 2–3 regions account for a median of 82% of PC1 variance (Supplementary Table S1). In roughly half of tracts (19/36), the top loadings include both positive and negative signs (Figure 4D), indicating that the primary axis of individual variation is not overall tract size but rather a trade-off between termination regions. For example, in the left arcuate fasciculus, posterior temporal-occipital regions load positively while the planum temporale loads negatively, capturing known individual variation in where the arcuate terminates along the temporal lobe. These mixed-sign loadings imply that reproducible individual differences often take the form of trade-offs between termination regions, information that would be lost under univariate analysis of individual region overlaps.

### 4.3 Practical implications for multi-site study design

Our results do not argue against diffusion scalars, which capture biologically meaningful microstructural properties. However, diffusion metrics carry the highest scanner-induced variance, and this gap persists after ComBat harmonization (harmonized diffusion ICC 0.58 remains below unharmonized shape ICC 0.69). For study designs where the scientific question can be addressed through geometric properties, shape metrics offer a more scanner-robust alternative without requiring harmonization.

For connectivity, not all regions contribute equally to reproducible individual signal. The per-tract loading structure reported here (Figure 4D, Supplementary Table S1) identifies which regions carry the most reliable between-subject variation, and these regions should be analyzed jointly to preserve the trade-off structure revealed by mixed-sign loadings. For studies using different pipelines or parcellations, a covariance-PCA on a small reference subset can identify the equivalent high-loading regions. This recommendation applies to study designs where individual differences are the target (e.g., group comparisons, biomarker development). For applications requiring the full connectivity representation, such as systems-level network modeling, population-stable connections with near-zero PC1 loadings remain essential as the structural scaffold of the circuit.

More broadly, multi-site studies may benefit from considering whether their scientific question requires voxel-level microstructural measures, or whether it can be addressed using higher-level derived quantities that carry less scanner-induced variance. Diffusion scalars remain essential when microstructural specificity is the object of study, but our results suggest that scanner-robust alternatives exist and may be preferable when the scientific question permits.

### 4.4 Limitations

The sample comprises 5 subjects across 4 scanners. Because each subject is scanned on multiple scanners, between-subject and within-subject (scanner-driven) variance can be separated, which is sufficient for the reproducibility measures reported here (ICC and subject discrimination). Estimating scanner sensitivity as a distinct quantity would require multiple sessions within the same scanner per subject, which was available for only 2 of 4 scanners, precluding reliable quantification. Future work could extend this framework using larger traveling-subject designs where within-scanner replication is available. Additionally, all analyses were conducted using a single tractography platform. While this could limit generalizability, it also ensures that cross-family comparisons are not confounded by differences between processing tools, as the platform used is one of few that provides tract-level shape, diffusion, and connectivity metrics within a unified pipeline.

## 5. Conclusion

Shape metrics achieve the highest baseline reproducibility across scanners. Connectivity metrics, under simple dimensionality reduction, yield subject-specific signatures that surpass all families at any level of harmonization. ComBat harmonization improves all families but does not eliminate the reproducibility hierarchy. These findings suggest that the question of which metrics to use in multi-site studies deserves at least as much attention as the question of how to harmonize them.

## Author Contributions

Sachit Anand: Formal Analysis, Software, Visualization, Writing (review and editing) Fang-Cheng Yeh: Software, Resources, Funding Acquisition, Writing (review and editing) Siva Venkadesh: Conceptualization, Methodology, Investigation, Formal Analysis, Software, Writing (original draft), Supervision

## Acknowledgments

Computational resources were provided by the Pittsburgh Supercomputing Center through NSF ACCESS allocations MED240021, MED230052 and CIS200026.

## Data and code availability

The MASiVar Cohort II dataset is publicly available on OpenNeuro (ds003416). Analysis and figure generation code is available at https://doi.org/10.5281/zenodo.19581307.

## Declaration of generative AI use

During the preparation of this work the authors used Claude (Anthropic) to assist with manuscript drafting, literature review, and organizational structuring. After using this tool, the authors reviewed and edited the content as needed and take full responsibility for the content of the published article.

