## Supplementary Figure S1 for "Reproducibility of Diffusion, Shape, and Connectivity Metrics Across Scanners: Implications for Multi-Site Tractography"

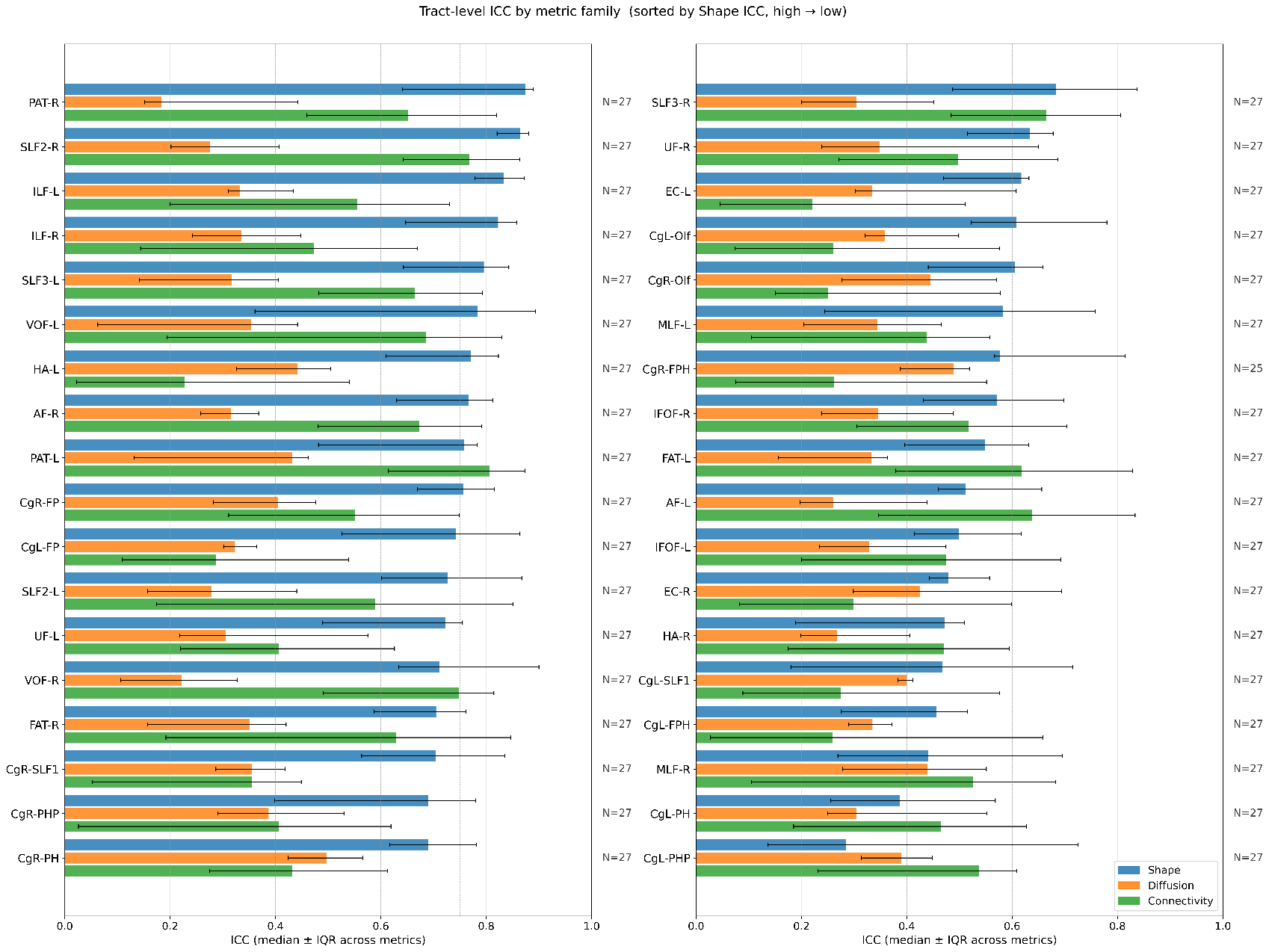

**Supplementary Figure S1. Tract-level ICC by metric family for all 36 association tracts.** Horizontal bar charts showing median ICC (with interquartile range across metrics) for shape (blue), diffusion (orange), and connectivity (green) within each tract. Tracts are sorted by descending shape ICC. N indicates the number of scanning sessions available per tract.
